# Insight into the structures of unusual base pairs in RNA complexes containing primer/template/adenosine ligand

**DOI:** 10.1101/2023.07.20.549964

**Authors:** Yuliya Dantsu, Ying Zhang, Wen Zhang

**Affiliations:** Department of Biochemistry and Molecular Biology, Indiana University School of Medicine, Indianapolis, IN 46202, USA

## Abstract

In prebiotic RNA world, the self-replication of RNA without enzymes can be achieved through the utilization of 2-aminoimidazole activated nucleotides as efficient substrates. The mechanism of RNA nonenzymatic polymerization has been extensively investigated biophysically and structurally by using the model of RNA primer/template complex which is bound by the imidazolium-bridged or triphosphate-bridged diguanosine intermediate. However, beyond the realm of guanosine substrate, the structural insight into how alternative activated nucleotides bind and interact with RNA primer/template complex remains unexplored, which is important for understanding the low reactivity of adenosine and uridine substrates in RNA primer extension, as well as its relationship with the structures. Here we use crystallography as method and determine a series of high-resolution structures of RNA primer/template complexes bound by ApppG, the close analog of dinucleotide intermediate containing adenosine and guanosine. The structures show that ApppG ligands bind to RNA template through both Watson-Crick and noncanonical base pairs, with the primer 3′-OH group far from the adjacent phosphorus atom of the incoming substrate. The structures indicate that, when adenosine is included in the imidazolium-bridged intermediate, the complexes are likely preorganized in a suboptimal conformation, making it difficult for the primer to in-line attack the substrate. Moreover, by cocrystallizing the RNA primer/template with chemically activated adenosine and guanosine monomers, we successfully observe the slow formation of the imidazolium-bridged intermediate (Ap-AI-pG) and the preorganized structure for RNA primer extension. Overall, our studies offer a structural explanation for the slow rate of RNA primer extension when using adenosine-5’-phosphoro-2-aminoimidazolide as a substrate during nonenzymatic polymerization.

## Introduction

According to RNA world hypothesis proposed by Crick^1^, Orgel^2^ and Woese^3^, RNA was the essential molecule to construct the first cell in prebiotic world with the functions of storing genetic information and catalyzing biochemical reactions^4, 5^. The idea of the RNA world hypothesis rationalizes how life began in its early stages, including the transition from simple molecules into basic RNA, and the later evolvement into complex ribozymes to catalyze primitive metabolism. For the emergence of RNA-based life, oligomerized RNA strands had to get replicated without enzymes^4, 6, 7^. For RNA’s self-replication, the building blocks or short chains need to be chemically activated, bind to the complementary template, arrange themselves in a specific conformation and get polymerization reaction occur to produce the new copy of RNA^8^. Considerable efforts have been dedicated over the years to the optimization of RNA nonenzymatic polymerization, like enhancing the fidelity and efficiency of the reaction, exploring alternative chemical activations, searching for effective catalysts, and augmenting the rate of polymerization^9^. One successful example is the discovery of derivatized-imidazole-activated ribonucleotides as reaction substrates, including nucleoside-5’-phosphoro-(2-aminoimidazolide)^10, 11^ and nucleoside-5’-phosphoro-(2-methylimidazolide)^12^, which can mediate the fast copying of RNA sequences in the presence of RNA primer, template, and divalent metal ion^10, 13^. The thermodynamics, kinetics and reaction mechanism of primer extension using this superior substrate has been extensively explored^14-17^. The nonenzymatic polymerization reaction is purely chemically catalyzed, and it requires the formation of a 5′-5′ phospho-imidazolium-phospho bridged dinucleotide intermediate^18^. The dinucleotide intermediate binds the RNA template with great affinity and well-defined conformation, including multiple inter- and intramolecular hydrogen bonds to preorganize the complex^19^. Consequently, the RNA primer extension proceeds stepwise with the 3′ hydroxyl group of the adjacent primer orientated to in-line attack the incoming phosphorus atom of the imidazolium-bridge and replace an activated nucleotide as the leaving group of nucleophilic reaction^18^.

To ensure the propagation of primitive life, the accurate transfer of the complete genetic information is critical^6, 7^. In the enzyme-free RNA replication, that will require the efficient incorporation of all four nucleotides and the replication of RNA strands containing mixed sequences. Generally, the error rate in nonenzymatic copying could easily lead to standstill of primer extension and the truncated information that is passed on from generation to generation^20^. The incorporation of G and C nucleotides occurs with higher rate and fidelity than A and U monomers^21-24^. The stalling effect arising from A and U monomers is possibly due to the weak binding of adenosine and uridine-containing substrates (including monomers or dinucleotide intermediates) to template, or the local geometry of RNA primer/template/A or U substrate complexes, which is less favorable for the nucleophilic attach of the 3′-OH group of RNA primer. Recently, the frequency of mismatches in RNA nonenzymatic polymerization and its relationship with dinucleotide intermediate mechanism has been investigated^25^, and the method of using deep-sequencing to identify mismatches in primer extension is developed^26^. Overall, a more in-depth understanding of how adenosine or uridine substrates affect the nonenzymatic polymerization is important and demands further exploration. Therefore, we decide to investigate the structural insights into the nonenzymatic primer extension when there is the adenosine substrate binding to RNA template and forming primer/template complex, asking if there are nonconventional binding motifs existing to affect the primer extension.

To understand how nucleotide substrates bind to RNA template and get preorganized for primer extension, many high-resolution crystal structures have been determined using the stable reaction substrate analogs, including the guanosine-5′-phosphoro-pyrazole monomer^27^ and the symmetrical 5′-5′ triphosphate-linked dinucleotide GpppG^28^. These studies indicate that the guanosine monomers bind the template through various modes of base pairing^29^. In contrast, as the molecular analog of imidazolium-bridged intermediate, GpppG was observed to bind to RNA template only through Watson-Crick base-pairs, with the 3′-hydroxyl of the primer positioned for nucleophilic attack^28^. Additionally, the time-resolved crystal structures also provided the atomic resolution insight into the binding motif of real activated substrate and the mechanism of nonenzymatic primer extension^19^. These structural observations suggest that the Watson-Crick base pairing between substrate and template result in a favorable preorganized geometry and high reactivity. Here we use the similar approach of crystallography and report the structural insight into the mechanism of nucleotide substrate binding and interacting with RNA when adenosine is present.

## Results and discussion

### Primer extension assays with adenosine-5’-phosphoro-(2-aminoimidazolide) (2-AIpA) substrate

We first ask whether the RNA primer can be nonenzymatically extended when the chemically activated adenosine monomer serves as the reaction substrate in solution (Figure 1A). We synthesize an RNA primer containing Cy3 fluorophore at its 5’ end (5’-Cy3-GUAGACUGACUGA-3’, Figure 1B), as well as different templates that contain uridine or cytidine residues at template to bind to 2-AIpA and 2-AIpG monomers. We then examine rates of RNA-templated primer extension by one nucleotide, when there are different activated nucleotides bound and incorporated at +1 and +2 positions. In the group of RNA primer/template system containing 3’-CC as template and 2-AIpG as substrate, primer extension proceeded quickly as expected, with an observed rate of 0.7 h^-1^. However, when the mixture of 2-AIpA and 2-AIpG was used as substrate in solution, with the 3’-UC or 3’-CU as template, the observed rates were dramatically reduced to 0.07 and 0.13 h^-1^, nearly 10-fold and 5-fold slower than the RNA extension with 3’-CC template (Figure 1C-E). In contrast, with the 3’-UU as template and 2-AIpA as substrate, the reaction became extremely slow (est. less than 0.01 h^-1^). Thus, our result suggests that once the A:U base pair is involved during primer extension, the reaction rate will significantly diminished. With the exclusive presence of A:U pair between template and substrate, and the polymerization can hardly proceed to the end of the template.

**Figure 1.**
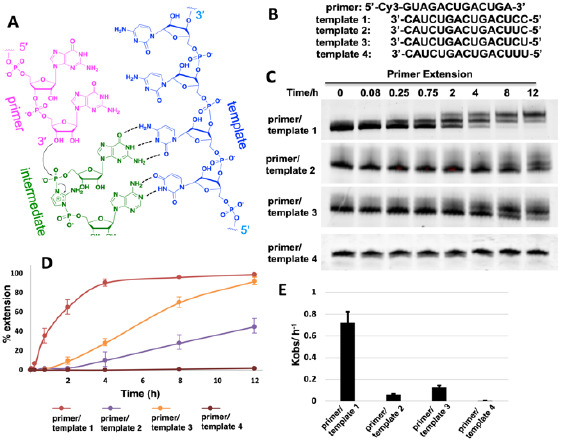
(A) Chemical representation of hypothetical template-directed polymerization with RNA primer, RNA template and 2-aminoimidazole bridged intermediate containing adenosine and guanosine (Ap-AI-pG). (B) Sequences of primer and templates used in the primer extension. (C) PAGE analysis of the reaction products over different times using different templates. Gels depicted are representative of triplicate experiments. (D) Comparison of primer extension with different templates. (E) Pseudo-first order rates of the reactions with different templates.

### Structures of the RNA primer-template complexes co-crystallized with AMP

We then decide to structurally explore how adenosine substrate binds to RNA template in nonenzymatic polymerization. We first cocrystallized AMP mononucleotide with self-complementary RNA oligonucleotide 5’-***TTAG***ACUUAAGUCU-3’, which is flanked on both ends by two locked thymidine nucleotides serving as the binding sites for AMP (Figure 2A). The locked nucleic acid (LNA) modification (denoted as bold and italic nucleotides) constrains the sugar into the 3’-endo conformation and helps the complex crystallization for high-resolution X-ray structures. The key crystallographic parameters are listed in Supporting Information. There is one RNA duplex/AMP complex per asymmetric unit. Like the structures determined previously^19^, the individual double helices are slip-stacked with one another end-to-end. Groups of three RNA duplex–ligand complexes form triangular prisms, and the central channel accommodates water molecules to bridge the neighboring complexes (Figure 2B).

**Figure 2.**
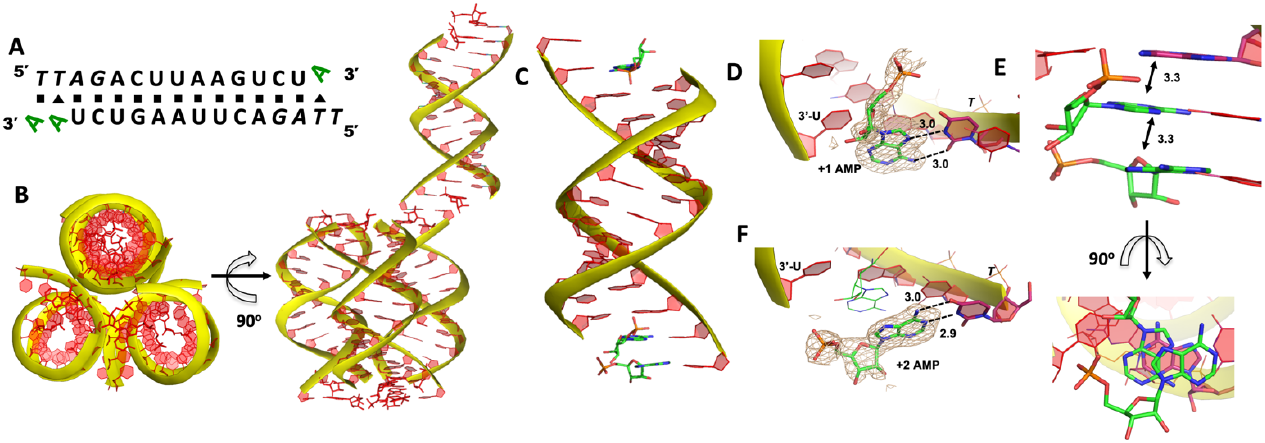
Crystal structure of RNA/AMP complex. (A) Diagram and designed duplex structure of RNA/AMP complex. AMP monomers (green) are bound at each end. Black square: Watson−Crick pairs. Black triangle: noncanonical base pairs. (B) The adjacent three RNA/AMP complexes assemble for crystal formation. The bound AMPs are stacked between two coaxial RNA duplexes. (C) Overall structure of RNA/AMP duplex. (D) The local structure of AMP binding to RNA template at +1 position. The corresponding *F*_*o*_ − *F*_*c*_ omit map contoured at 1.5σ (wheat mesh) indicates the noncanonical base pair. (E) The bound AMP at +1 position is well stacked with AMP at +2 position and upstream adenine. (F) The local structure of AMP binding to RNA template at +2 position, forming the Watson-Crick base pair. All the significant distances are labelled.

At one end of the duplex, there are two AMP monomers bound to the templating locked thymidines consecutively (Figure 2C). At +1 position, a noncanonical A:T base pair was observed, mediated by two hydrogen bonds: the N7 of the adenine was 3.0 Å from the N3 of templating thymine, and the exocyclic amine of the AMP was 3.0 Å from the exocyclic oxygen atom of thymine (Figure 2D). Interestingly, by forming this unique base pairing, the adenine nucleobase at +1 position was sandwiched between the AMP monomer bound at +2 position and the adenine nucleobase which is located at the templating strand and adjacent to the templating thymidine (Figure 2E). The internucleobase distance between the two bound adenine bases is approximately 3.3 Å based on analysis using the CONTACT program from CCP4^30^, and the distance between the adenine bound at +1 position and “upstream” adenine is also 3.3 Å. This binding motif of AMP is different from what was observed in the structures of template/primer/guanosine substrate complexes, and the distance between the 3′-OH group of primer and the phosphorus atom of bound AMP was over 10 Å. This local structure may explain the slow primer extension rate when using adenine monomer as substrate. At +2 position, the AMP is forming Watson−Crick base pair with the thymine nucleobase (hydrogen-bond lengths, 3.0 and 2.9 Å, Figure 2F). Unfortunately, the sugar moieties of both bound AMP monomers are not ordered enough to predict the sugar conformation. At the other end of the duplex, only one AMP monomer was observed to bind the template at +1 position via the noncanonical base pairing. The overall binding motif remains the same as the other +1 bound AMP.

To explore whether the noncanonical A:T pair widely exist when binding to RNA primer/template, we next cocrystallized another self-complementary RNA 5’-***T***^***m***^***C***^***m***^***CG***ACUUAAGUCG-3’ with both AMP and GMP monomers to bind the two nucleotides overhang 5’-***T***^***m***^***C*** at both ends (Figure 3A, ^***m***^***C*** represents the locked 5-methyl cytidine residue). The structure was determined to 1.45 Å resolution by molecular replacement using the previously determined RNA duplex as a search model^19^. As in the structures with the similar sequences, the individual double helices are slip-stacked with one another end-to-end. At one end, one AMP and one GMP nucleotides are clearly bound to the RNA, fully occupying the two-nucleotide binding sites. The adenine and guanine nucleobases of both monomers were defined with clear electron densities (Figure 3B), while the sugars and monophosphate groups were highly disordered. At +1 position of the primer/template complex, the guanine nucleobase of GMP was forming Watson-Crick base pair with ***mC*** LNA nucleotide (hydrogen-bond distances, 3.1, 3.0 and 2.8 Å, Figure 3C). At +2 position, the unusual base-recognition motif was observed again, via the previously described noncanonical A:T interactions (hydrogen-bond distances, 2.7 and 3.1 Å, Figure 3D). Overall, B factor and electron density fitting indicated that the bound GMP monomer at +1 position was more structurally ordered than AMP bound at +2 position, which is consistent with the previous structural and biophysical data^27, 31^, that the downstream bound substrate enhances the binding affinity of upstream monomer.

**Figure 3.**
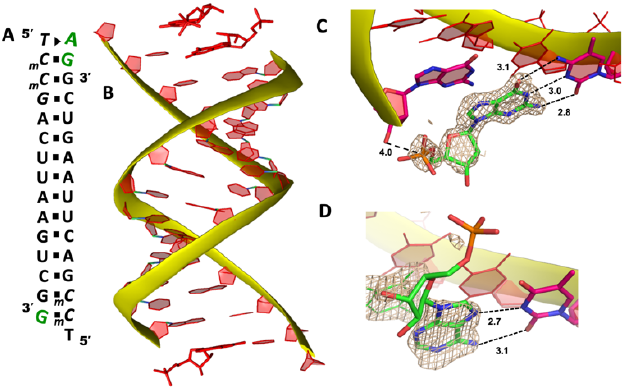
The structure of RNA/AMP/GMP complex. (A) Diagram and designed duplex structure of RNA/AMP/GMP complex. AMP and GMP monomers (green) are bound at each end. Black square: Watson-Crick base pairs. Black triangle: noncanonical base pair. (B) Overall structure of RNA/AMP/GMP duplex. (C) The local structure of GMP binding to RNA template at +1 position. The corresponding *F*_*o*_ − *F*_*c*_ omit map contoured at 1.5σ (wheat mesh) indicates the Watson-Crick base pair. (D) The local structure of AMP binding to RNA template at +2 position, forming the noncanonical base pair. All the significant distances are labelled.

It is noteworthy that, the unique A:T base pair was observed in both structures of RNA/mononucleotides complexes. By forming the unusual base pair, the sugar and phosphate groups of AMP point to the spacious major groove of helix and avoid the potential steric hindrance with the neighbouring bound monomer. Similar binding motif was also observed when two guanosine monomers bound to the consecutive cytosine templates^27^. However, in previously determined structures, half of the monomer-template binding motifs were consecutive Watson-Crick base pairing, which is different from AMP-template binding. According to the current two structures, it’s more frequent to observe AMP forming the noncanonical base pair with the templating thymidine when another neighbouring monomer is at presence. After forming the noncanonical base pair, the distance between the phosphate groups of two bound monomers is much longer (over 10 Å) than usual, which makes it difficult for the two monomers to have nucleophilic interaction between each other. Considering this information, we propose that in RNA nonenzymatic polymerization using chemically activated adenosine as a substrate, there is a higher likelihood of forming the noncanonical base pair between the substrate and template. This, in turn, inhibits the formation of the imidazolium-bridged dinucleotide substrate on the template. This structure highly disfavours the RNA primer extension, which probably explains the slow reaction rate when using 2-AIpA as substrate.

### Structure of GpppA bound to RNA template

Our RNA/AMP complex structures indicate that, when the activated adenosine monomer is included in the polymerization reaction and bound to template, formation of imidazolium-bridged intermediate on template will be challenging. Next, we explore whether the adenosine-containing dinucleotide intermediate, which is a product assumably formed in the solution, can indeed bind to the RNA template in a favorable conformation for the primer extension process. To study the complex structures when the imidazolium-bridged dinucleotide substrate binds to RNA template, the *P*^*1*^,*P*^*3*^-diguanosine-5′-triphosphate (GpppG) had been used as close analog and cocrystallized with RNA primer/template complex to interpretate the preorganization and reactivity of dinucleotide intermediate^28^. Therefore, we decide to use the commercially available *P*^*1*^-5′-adonesine-*P*^*3*^-5′-guanosine triphosphate (ApppG) to mimic the imidazolium-bridged intermediate (Ap-Im-pG) in nonenzymatic polymerization, and structurally study how the dinucleotide binds to RNA template via Watson-Crick or noncanonical base pairs.

We first cocrystallized ApppG with RNA 5’-***T***_***m***_***C***_***m***_***CG***ACUUAAGUCG-3’, where the 5’-***T***_***m***_***C*** overhang served as the binding site for ApppG (Figure 4A). The RNA crystallized with ApppG with the same symmetry as in the RNA-AMP complexes. The space group is P3_1_21, and there is one RNA duplex with two bound ApppG molecules per asymmetric unit. At each end of the RNA duplex, the overhanging 5’-***T***_***m***_***C*** binding site is fully occupied by the dinucleotides (Figure 4B). As in the RNA–AMP complex structures, the RNA double helices are A-form and the duplexes slip-stack on each other to form extended columns. At one end, the ApppG ligand forms two Watson-Crick base pairs with the 5’-***T***_***m***_***C*** overhang (Figure 4C, hydrogen bonds distances 2.8-3.1 Å). The guanine and adenine nucleobases are coplanar and stacked with the upstream primer and the neighboring duplex. The electron density suggests that the ribose sugar of guanosine at +1 position is in the C3′-*endo* A-form conformation, while the sugar of adenosine is disordered at +2 position. This observation is similar to the structure of RNA/GMP/AMP complex, which is consistent with the conclusion that the downstream bound substrate enhances the binding the sandwiched monomer. The electron density indicates that, the triphosphate group of ApppG dinucleotide exhibits a moderate level of order, enabling us to determine its geometric properties.

**Figure 4.**
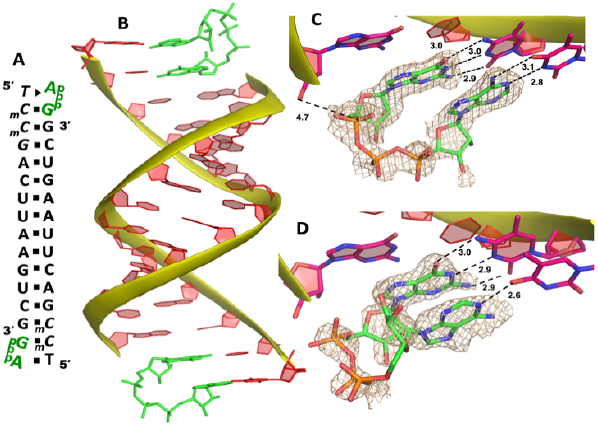
The structure of RNA/ApppG complex. (A) Diagram and designed duplex structure of RNA/ApppG complex. ApppG ligand (green) is bound at each end. Black square: Watson−Crick pairs. Black triangle: noncanonical base pairs. (B) Overall structure of RNA/ApppG duplex. (C) At one end, ApppG forms two Watson-Crick base pair with template with the moderately ordered linkage. (D) At the other end of duplex, ApppG is forming one Watson-Crick and one noncanonical base pair with template. The wheat color mesh indicates the corresponding *F*_*o*_ − *F*_*c*_ omit map of ApppG contoured at 1.5σ. All the significant distances are labelled.

As the close analog of the imidazolium-bridged intermediate, it’s important that the triphosphate linkage of ApppG is preorganized in a favorable conformation for the SN2 primer extension reaction. The distance between the primer 3′-hydroxyl and the phosphorus atom of the closest phosphate of ApppG is 4.7 Å, and the angle between the 3′-OH and the bridging P-O bond of GpppG is 110° (Figure 4C). This distance is significantly longer than what we observed in RNA-GpppG structure (4.1 Å)^28^. Unlike the RNA-GpppG structure that a Mg^2+^ ion was coordinating the triphosphate linkage in a well-defined conformation, no metal ion is observed surrounding the triphosphate group of ApppG and the reaction angle for the in-line attack by the primer is less favorable (110° vs 126°). Overall, the local structure of ApppG binding to RNA template suggests that the dinucleotide intermediate containing adenosine nucleotide is unlikely to be optimal for the RNA primer extension.

At the other end of the duplex, ApppG binds to the template in a distinctly different manner (Figure 4D). The guanosine adjacent to the primer is Watson–Crick base paired with the template _***m***_***C*** through three hydrogen bonds. However, the adenosine nucleobase forms a noncanonical A:T “pair”. The adenine nucleobase is well stacked with the upstream guanine, and doesn’t form any hydrogen bond with the templating thymine. However, the distance between N1 of adenosine and O4 of ***T*** residue is measured to be 2.6 Å, close enough to form a hydrogen bonding. Considering the potential tautomeric equilibrium in adenosine and the barely fitted electron density on the nucleobase, it is possible that the adenosine forms a weak base pair with the templating ***T*** by one hydrogen bond. The triphosphate linkage and sugar of adenosine are highly disordered, making it difficult to define the geometry of the linkage and how it benefits the SN2 primer attach. The overall structure suggests that ApppG likely binds to the RNA template with a weaker affinity than GpppG, and the primer/template/ApppG complex is unlikely to be structurally optimal for the primer extension.

To explore whether the downstream bound ligand can enhance the binding of adenine-containing intermediate and better preorganize the adenosine nucleotide, we then cocrystallized ApppG with another RNA 14mer, 5’-_***m***_***CT***_***m***_***CG***ACUUAAGUCG-3’, where the locked thymidine is adjacent to the primer and serves as the binding site of adenosine (Figure 5A). The structure was determined to 1.95 Å, and the crystal grew with the same symmetry as other RNA-ligand complexes. The overall crystal structure remains the same molecular packing patterns, and there is one ApppG dinucleotide ligand binding to 5’-_***m***_***CT*** template at each end (Figure 5B). However, with the different template, the binding mode of ApppG is different from that of 5’-***T***_***m***_***C*** template. At both ends, the electron density of adenine and guanine nucleobases strongly indicates that the ligand binds to template only via Watson-Crick base pairing and the hydrogen bonds range from 2.8 to 3.1 Å (Figure 5C). At both ends, the triphosphate linkage is moderately ordered, and the distance between the 3′-hydroxyl group of the primer and the phosphorus center of triphosphate linkage is measured to be 4.7 Å, also significantly longer than the distance in RNA-GpppG complex. Interestingly, one important water molecule is observed close to ApppG (Figure 5D). This well-ordered water molecule is forming two hydrogen bonds with N7 of adenosine (2.7 Å) and one of the nonbridging oxygens of the triphosphate linkage (3.0 Å), which is likely to contribute to stabilize ApppG. Based on our structural analysis, we suggest that the downstream G:C base pair plays a crucial role in impeding the formation of noncanonical A:T pairs. Instead, it enforces the A:T Watson-Crick base pair at the +1 position of the primer extension. Nevertheless, it’s worth noting that the RNA-ApppG complex typically adopts a suboptimal conformation for the SN2 nucleophilic attack, despite this beneficial effect of the G:C base pair.

**Figure 5.**
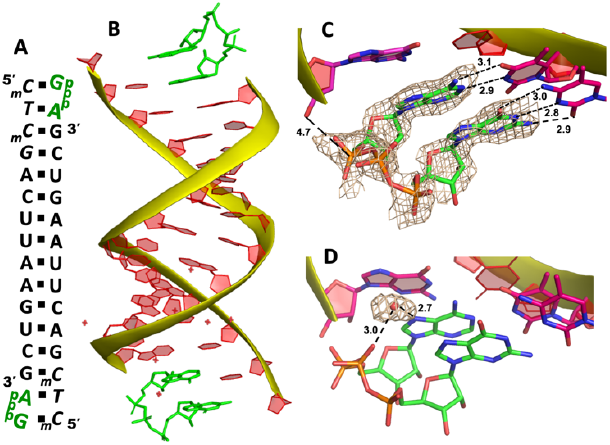
The structure of RNA/ApppG complex. (A) Diagram and designed duplex structure of RNA/ApppG complex. ApppG ligand (green) is bound at each end. Black square: Watson−Crick pairs. (B) Overall structure of RNA/ApppG duplex. (C) At both ends, ApppG forms two Watson-Crick base pairs with template with the moderately ordered linkage. (D) One water molecule is forming two hydrogen bonds to bridge the triphosphate linkage and adenosine nucleobase. The wheat color mesh indicates the corresponding *F*_*o*_ − *F*_*c*_ omit map of ApppG contoured at 1.5σ. All the significant distances are labelled.

### Structure of RNA-GpppA containing A:C mismatch

In replication, maintenance of fidelity in genetic information transfer is critical for the accurate replication of life. In the modern world, the genetic molecule of DNA undergoes replication with remarkable precision, thanks to the regulation of DNA polymerase. This enzyme is capable of discerning and correcting mismatched base pairs through proofreading mechanisms^32^.

In comparison, the nonenzymatic polymerization is error-prone, because it is lack of mismatch-repairing enzymes and the fidelity entirely depends on base pairing. The error rate in nonenzymatic copying could easily lead to standstill of primer extension and the truncated/mutated genetic information^20^. Therefore, it is critical to understand the origin and chemistry of mismatches in prebiotic nonenzymatic copying.

In this study, we decide to explore the possibility of A:C mispair in nonenzymatic polymerization. In natural biology, DNA is able to tautomerize via proton transfer to form irregular A:C mispair^33, 34^, which is important for the origin of spontaneous point mutations. A:C mismatched pair is stabilized by three hydrogen bonds, and it’s believed to play an important role in the gene replication errors^35^. To investigate whether the A:C mismatch could occur and affect the fidelity of RNA enzyme-free replication, we decided to perform structural studies on ApppG dinucleotide when it binds to RNA template through mismatched base pair.

We cocrystallized ApppG ligand with RNA 5’-_***m***_***C***_***m***_***C***_***m***_***CG***ACUUAAGUCG-3’, in which two locked methylcytidine served as binding sites (Figure 6A). The complex successfully crystallized and the structure was determined to 1.5 Å. There is one ApppG dinucleotide ligand binding to 5’-_***m***_***C***_***m***_***C*** template at each end (Figure 6B). In the structure, ApppG binds to the template through unusual conformational motif. At +1 position, guanine nucleobase is binding to the templating _***m***_***C*** via Watson-Crick base pair (3.2, 2.9, 2.8 Å), and the ribose sugar is in 3′-*endo* conformation. At +2 position, the electron density indicates that the adenine nucleobase is well stacked with the guanine base at +1 position, and the adenine nucleobase is forming only one hydrogen bond with _***m***_***C* (**3.1 Å, between N3 of adenine and N4 of _***m***_***C***). The sugar of adenosine is highly disordered. The electron density associated with the triphosphate linkage is ordered, and the distance between the 3′-hydroxyl group of the primer and the adjacent phosphorus center is about 4.6 Å (Figure 6C). The overall structure indicates that when there are mismatched nucleotides in the RNA template, the adenosine-containing dinucleotide intermediate is prone to forming a noncanonical weak base pair. This local complex is then stabilized by stacking interactions, resulting in a suboptimal structure for the primer extension reaction.

**Figure 6.**
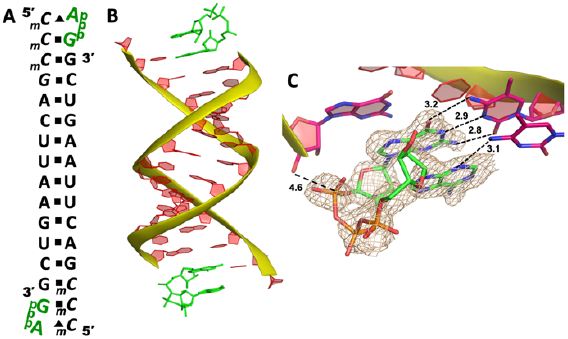
The structure of RNA/ApppG complex containing A:C mismatch. (A) Diagram and designed duplex structure of RNA/ApppG complex. ApppG ligand (green) is bound at each end. Black square: Watson−Crick pairs. Black triangle: noncanonical base pairs. (B) Overall structure of RNA/ApppG duplex. (C) At both ends, ApppG forms one Watson-Crick and one noncanonical base pairs with template with the moderately ordered linkage. The wheat color mesh indicates the corresponding *F*_*o*_ − *F*_*c*_ omit map of ApppG contoured at 1.5σ. All the significant distances are labelled.

### Structure of imidazolium-bridged adenosine, guanosine intermediate (Ap-AI-pG) bound to RNA template

It has been previously shown that nonenzymatic RNA primer extension is mediated by a highly reactive imidazolium-bridged dinucleotide intermediate, which binds to template with great affinity and well-preorganized conformation to benefit the nucleophilic attack of the primer^19, 36^. We therefore attempt to explore whether the dinucleotide intermediate containing different nucleobases can be formed and how this intermediate binds to RNA primer/template complex in nonenzymatic polymerization.

We first followed the previously reported method in time-resolved structure determination^19^, by cocrystallizing GMP and AMP monomers with RNA 5’-***T***_***m***_***C***_***m***_***CG***ACUUAAGUCG-3’ to form the primer/template/monomers complex (Figure 7A). After obtaining the crystals, we tried to exchange the inactivated nucleotides with the activated guanosine-5’-phosphoro-2-aminoimidazolide (2-AIpG) and adenosine-5’-phosphoro-2-aminoimidazolide (2-AIpA) by soaking the crystals in a solution of activated monomers. By soaking the crystals for different periods, we sought to observe the formation of imidazolium-bridged Ap-AI-pG intermediate on the 5’-***T***_***m***_***C*** binding site. However, even after screening a wide range of soaking times (up to 10 days) and activated monomer concentrations (up to 50 mM), we didn’t observe the formation of dinucleotide intermediate in the crystal structures. The sugar and phosphate moieties of both monomers are highly disordered, so that it’s impossible to conclude whether the bound ligands are 2-aminoimidazole-activated monomers or the nucleoside-5’-monophosphate molecules.

**Figure 7.**
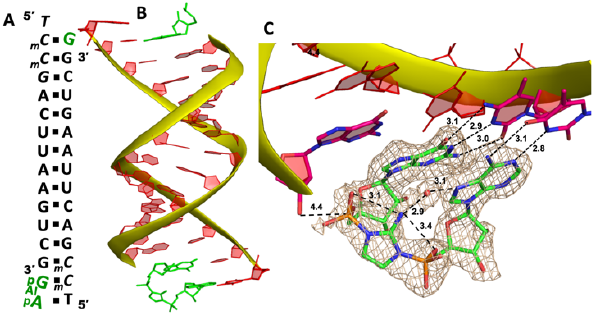
The structure of RNA/Ap-AI-pG complex. (A) Diagram and designed duplex structure of RNA/Ap-AI-pG complex. Ap-AI-pG ligand (green) are bound at one end. Black square: Watson−Crick pairs. (B) Overall structure of RNA/Ap-AI-pG duplex. (C) At one end, Ap-AI-pG is formed and binding the template via two Watson-Crick base pairs. One water molecule is forming two hydrogen bonds to bridge the 2-aminoimidazole linkage and adenosine nucleobase. The wheat color mesh indicates the corresponding *F*_*o*_ − *F*_*c*_ omit map of ApppG contoured at 1.5σ. All the significant distances are labelled.

To structurally visualize the RNA/Ap-AI-pG complex, we then decided to cocrystallize RNA with 2-AIpG and 2-AIpA monomers. In the crystallization drop, the activated monomers ought to bind to the corresponding RNA template when we set up the crystallization. We expect that, during crystal growth, intermolecular interaction between 2AIpG and 2AIpA could occur on template to form the intermediate, or the slowly emerged imidazolium-bridged dimer in solution can exchange with 2-AIpG and 2-AIpA monomers bound to the template. It took about 40 days from setting up crystallization to the diffraction data collection. The optimal structure was determined to 1.7 Å, and there are two RNA/ligands complexes observed in one asymmetric unit. At one end of the A-helical duplex, we observe electron density consistent with imidazolium-bridged Ap-AI-pG bound to the template (Figure 7B). Both guanosine and adenosine are well ordered and form canonical Watson-Crick base pairs with the templating _***m***_***C*** and ***T*** residues (hydrogen bonds from 2.8 to 3.1 Å, Figure 7C). Both ribose sugars are restrained to 3′-*endo* conformations. Critically, the electron density between the two nucleotides agrees with the formation of an imidazolium bridge, and the ordered structure enable us to estimate the SN2 attack geometry. The distance between the 3′-OH of the primer and the incoming P atom of Ap-AI-pG molecule is 4.4 Å. This distance is shorter than all the distances we observed in RNA/ApppG complex structures where noncanonical A:T mismatch is present. It is even slightly shorter than the distance observed in the RNA complex containing diguanosine intermediate Gp-AI-pG (4.6 Å)^19^. Furthermore, the important O-P-N angle of primer attack is 122.4°, which is less than the previously observed angles of 132° and 170° in RNA/Gp-AI-pG structure. The phosphate-aminoimidazole-phosphate bridge is stabilized by hydrogen bonds between the 2-amino group of the imidazolium moiety and the non-bridging oxygen atoms of the flanking phosphates (3.1 and 3.4 Å), and this unique geometry was also observed in Gp-AI-pG structure. In the structure, we also clearly observe electron density near the N7 of the adenine nucleobase that appears to represent a water molecule. This water molecule is located in the major groove of the RNA duplex, and forms two hydrogen bonds with N7 atom of adenine and 2-amino group of the imidazolium-bridge (distances 3.1 and 2.9 Å). This proposed water mediates the intramolecular hydrogen bonds and bridges the imidazolium linkage and nucleobase. It potentially stabilizes the weakly bound adenosine ligand and preorganizes the structure of primer/template/intermediate complex for RNA polymerization. At the other end of the duplex, only one guanosine monomer is observed bound to _***m***_***C*** template, and the Ap-AI-pG dinucleotide is not formed. Table 1 summarizes all the RNA/ligand structures, for comparing the important conformational parameters for primer extension.

**Table 1.** Comparison of the key conformational parameters in difference crystal structures of RNA/substrate complexes. na=not available.

| Entry | Template | Bound ligands | Base pair at +1 position | Base pair at +2 position | distance | angle | Additional interaction |
| --- | --- | --- | --- | --- | --- | --- | --- |
| 1 | 5'-*TT* | AMP | noncanonical | Watson-Crick | > 10 | n.d. | na |
| 2 | 5'-*T<sub>m</sub>C* | AMP/GMP | Watson-Crick | noncanonical | 4.0 | n.d. | na |
| 3 | 5'-*T<sub>m</sub>C* | ApppG | Watson-Crick | Watson-Crick /noncanonical | 4.7 | 110° | na |
| 4 | 5'-*<sub>m</sub>CT* | ApppG | Watson-Crick | Watson-Crick | 4.7 | 131° | water mediated bridge |
| 5 | 5'-*CC* | ApppG | Watson-Crick | noncanonical | 4.6 | 125° | na |
| 6 | 5'-*T<sub>m</sub>C* | Ap-AI-pG | Watson-Crick | Watson-Crick | 4.4 | 122.4° | water mediated bridge |

## Discussion

In modern biology, during the replication of genetic information, the Watson-Crick base pairing geometry plays a dominant role in DNA polymerization. This is particularly significant due to the numerous hydrophobic, electrostatic, and hydrogen-bonding interactions that facilitate the discrimination of mismatched pairs. This recognition motif, evolved over time, serves as the foundation for precise and efficient transfer of genetic information. In contrast, in the prebiotic world, nonenzymatic gene replication lacks the constraints imposed by enzymes to aid in substrate/template recognition. As a result, a wide range of base pairing possibilities is allowed, enabling the full utilization of the hydrogen bond donor and acceptor groups on the nucleobases. The previous study suggest that the nonconventional base pairing motifs are possible to occur between the activated guanosine monomer and cytidine template during nonenzymatic RNA copying in solution and in crystals^27^. In the current work, we observe that adenosine substrate can also form various unusual base pairs with the thymidine template under nonenzymatic conditions. Moreover, in specific structures, the A:T base pair exhibits exceptional weakness, relying only on a single hydrogen bond and the stacking interaction between neighboring nucleobases. Indeed, if such weak A:T base pairs occur during RNA polymerization in solution, the process with the adenosine substrate could lead to errors and significant stalling effects during primer extension. This phenomenon explains the reduction in the polymerization rate observed when we attempt to extend the primer in the presence of adenosine at the +1 or +2 position. To overcome the problem, one possible strategy in RNA world could be the evolvement of G/C-rich ribozymes which can enforce the A:U Watson-Crick base pair and thereby significantly enhance the RNA template replication accuracy and efficiency, thus expanding the genetic code in primitive RNA genome.

The RNA enzyme-free polymerization relies on the involvement of the imidazolium-bridged dinucleotide intermediate, which can bind the RNA template with great affinity and interact with the RNA primer with high reactivity. Through the structural analysis of the actual substrates Gp-AI-pG and its closely related analog GpppG, compelling evidence has been obtained. The high-resolution structures demonstrate that the dinucleotide intermediate, particularly the imidazolium-bridge, plays a crucial role in preorganizing the primer/template/substrate complex, which significantly enhances the reaction rate^19^. Many inter- and intramolecular hydrogen bonds contribute to stabilize the structure. If a similar mechanism occurs with the imidazolium-bridged G, A-dinucleotide substrate, a question arises: why does the presence of the adenosine residue result in such a significant retardation effect during primer extension? In our structure, the only difference when replacing Gp-AI-pG intermediate with Ap-AI-pG is the loss of one hydrogen bond when binding to template (A:U vs G:C). Ap-AI-pG intermediate can still form 5 hydrogen bonds with template via Watson-Crick base pairs, and the hydrogen bonds between the 2-amino group of the imidazolium bridge and the flanking non-bridging phosphate oxygen atoms remain. The distance between the 3′-OH of the primer and the incoming P atom of Ap-AI-pG is also comparable to that in RNA/Gp-AI-pG structure (4.4 Å vs 4.6 Å). However, observing the formation of the imidazolium bridge in Ap-AI-pG takes significantly more time compared to Gp-AI-pG. One potential explanation is that, the dinucleotide intermediate is formed after the mononucletide binds the pairing template. Frequent noncanonical base pairing occurs when the 2-AIpA monomer binds to the uridine residue on the template, which makes the intermolecular interaction between 2-AIpA and 2-AIpG extremely challenging (distance between the phosphorus atoms > 10 Å). The Ap-AI-pG intermediate we observed was the product when both 2-AIpA and 2-AIpG were bound via Watson-Crick base pairing, which is uncommon.

In addition, our RNA/ApppG structures exhibit the presence of multiple A:T base pairing motifs, which serve as substitutes for the weak A:T Watson-Crick base pair. Once adenosine is forming the noncanonical base pair, the linkage between the nucleotides shifts towards the major groove of the duplex. As a result, the distance critical for the primer attack to occur increases. In contrast, in all the structures of RNA/GpppG and RNA/Gp-AI-pG, the dinucleotide ligands bind to the template exclusively through Watson-Crick base pairs, leading to a preorganized local geometry necessary for the SN2 in-line attack. Based on the obtained structural information, there is another possibility to consider. If the adenosine-containing intermediate is formed in solution without the aid of an RNA template, it could adopt multiple binding motifs simultaneously when approaching and binding to the template. Among the various motifs, only the Watson-Crick fashion possesses the favorable structure for primer extension. However, this motif faces competition from numerous unfavorable structures at the binding site, creating a challenging environment for primer extension. It eventually leads to the slow reaction rate.

It is also important to note that in our primer extension reactions, there is a possibility of alternative mismatched base pairs forming. For example, the guanosine nucleobase has the ability to form a wobble pair with a uridine residue on the template. This means that the Gp-AI-pG intermediate could potentially bind to the CU template, which might be one of the reasons for the slow rate observed in our primer extension experiments. All the structures presented here only represent a single instance of the primer/template/intermediate complex in the presence of an adenosine substrate in solution. To address the questions regarding the structural binding of the Gp-AI-pG intermediate to the CU template, as well as the competitive nature between wobble-paired Gp-AI-pG and Watson-Crick-paired Ap-AI-pG substrate, further investigation in the future is required.

## Materials and methods

All the oligonucleotides were chemically synthesized in a 1.0 µmol scale using an ABI394 Synthesizer. The RNA and locked nucleotide phosphoramidites used in this work were purchased from Glen Research and ChemGenes Corporation. All the oligonucleotides were prepared with DMTr-on, and in-house deprotected using concentrated ammonium hydroxide solution for 16h at 55 °C. The RNA strands were additionally desilylated with Et_3_N•3HF solution to remove TBDMS groups. The 5’-DMTr deprotection was either carried on using the commercial Glen-Pak purification cartridge (Glen Research Inc.), or performed by adding 3% trichloroacetic acid solution, followed by neutralization to pH 7.0 with 1 M TEAAc buffer. The oligonucleotides were purified by reversed phase HPLC on a Hamilton PRP-C18 HPLC Column (250 × 21.2 mm) at a flow rate of 10 ml/min. The oligonucleotides were collected, lyophilized, redissolved in water, and concentrated as appropriate for downstream experiments. Data were collected at the LS-CAT beamlines at Argonne National Laboratory. Datasets were processed using HKL2000 and DENZO/SCALEPACK^37^. All structures were solved by molecular replacement. The refinement protocol includes simulated annealing, positional refinement, restrained B-factor refinement, and bulk solvent correction^38^. The topologies and parameters for locked nucleotides _***m***_***C***(LCC), ***G***(LCG), ***T***(LNT), GpppA (GA3), and dinucleotide intermediate (GMA) were constructed and applied. Detailed experimental protocols are provided in Supporting Information.

## Acknowledgements

We thank the Zhang lab for helpful discussions, insightful commentary and careful revision of the manuscript. We thank Dr. Zdzislaw Wawrzak from the Life Sciences Collaborative Access Team beamline 21-ID-D at the Advanced Photon Source, Argonne National Laboratory (USA). This research used resources of the Advanced Photon Source, a U.S. Department of Energy (DOE) Office of Science User Facility operated for the DOE Office of Science by Argonne National Laboratory under Contract No. DE-AC02-06CH11357. Use of the LS-CAT Sector 21 was supported by the Michigan Economic Development Corporation and the Michigan Technology Tri-Corridor (Grant 085P1000817).

## Accession numbers

Atomic coordinates and structure factors for the reported crystal structures have been deposited with the Protein Data bank under accession code 8SXL, 8SX6, 8SWO, 8SX5, 8SWG, 8SY1.

