## supporting information for "Insight into the structures of unusual base pairs in RNA complexes containing primer/template/adenosine ligand"

### CONTENTS

1. General Methods S2

2. X-ray Crystallographic Studies S3

### 1. General Methods

**1.1 General Information.** All chemicals were purchased from Fisher Scientific, VWR or Sigma-Aldrich (St. Louis, MO). 2-aminoimidazole HCl was purchased from CombiBlocks, Inc. (San Diego, CA). All the nucleoside phosphoramidites for solid-phase synthesis were from Chemgenes Corporation or Glen Research. All the screening buffers for crystallization are from Hampton Research and Jena Bioscience. The syntheses of Adenosine-5′-*O*-phosphoro-(2-aminoimidazole) (2-AIpA) and Guanosine-5′-*O*-phosphoro-(2-aminoimidazole) (2-AIpG) followed the reported protocol[1].

#### 1.2 Determination of RNA concentration

Concentrations of the aqueous RNA samples were determined by their UV absorption at 260 nm on a Thermo Scientific Nanodrop 2000c Spectrophotometer (Waltham, MA). The theoretical molar extinction coefficients of these samples at 260 nm were provided by Integrated DNA Technologies, Inc..

#### 1.3 RNA primer extension

RNA primer and template were pre-mixed in 20 µL of a solution containing 200 mM HEPES buffer, pH 8.0, 1 uM primer and 5 uM template and 50 mM $MgCl_2$. The primer extension reaction was initiated by adding the activated monomers (2-AIpA and 2-AIpG) from 100 mM stock solutions to a final concentration of 10 mM in the reaction. At the indicated time points, 2 µL aliquots were taken from the reaction and quenched with 28 µL of 90 % (v/v) aqueous formamide containing 15 mM EDTA. Samples were heated at 95 °C for 2 min, cooled to room temperature, and analysed on 18 % (19:1) denaturing PAGE gels containing 7 M urea. Gels were imaged using a Bio-Rad ChemiDoc™ Imager. The resulting gel images were analysed using ImageJ software. The reactions were repeated by triplicate experiments.

#### 1.4 Crystallization

The locked nucleotides-containing RNA samples (0.5 mM duplex) were mixed with different binding ligands (20 mM), heated to 90 °C for 2 min, then slowly cooled to 4 °C prior to crystallization. The sitting drop vapor diffusion crystallization screening was performed using an Oryx4 Douglas Instruments robotic system. Screening conditions included Nucleic Acid Mini Screen Kits, Natrix HT, Index (Hampton Research) and Nuc-Pro-HTS (Jena Bioscience). Crystals grew at 18 °C in the vibration free IN45 Chilling/Heating Incubator (Torrey Pines Scientific).

**1.5 Data collection, structure determination and refinement.** Crystals were mounted using CrystalCap™ SPINE HT with CryoLoops (Hampton Research). Crystals were soaked for 2 min in a drop of mother liquor solution which contained 30% glycerol as a cryo-protectant, before dipped into liquid nitrogen.

Diffraction data were collected at a wavelength of 1 Å under a liquid nitrogen stream at -174 °C at the Life Sciences Collaborative Access Team (LS-CAT) beamline 21-ID-D at the Advanced Photon Source, Argonne National Laboratory (USA). The crystals were exposed for 0.1 s per image with a 0.2° oscillation angle. The distances between detector and the crystal were set to 100-300 mm. Raw diffraction data was processed using HKL2000 and DENZO/SCALEPACK. The structure was determined by molecular replacement using the published structure (PDB code 6c8I) as search model. The structure was refined using Refmac[2] in the CCP4 suite. The refinement protocol included simulated annealing, positional refinement, restrained B-factor refinement, and bulk solvent correction. The topologies and parameters for synthetic  for locked nucleotides $_m$***C***(LCC), ***G***(LCG), ***T***(LNT), GpppA (GA3), and dinucleotide intermediate (GMA) were created using ProDrg[3] in CCP4 suite and applied during refinement. After several cycles of refinement, a number of ordered waters and metals were added.

**2. X-ray Crystallographic Studies.** Optimal crystallization conditions, data collection, phasing, and refinement statistics of the determined structures are listed in Tables S1, S2 and S3.

**Table S1. Optimized conditions for crystallization of RNA-monomer complexes**

| entry | Binding template | ligands | Optimized crystallization conditions |
|---|---|---|---|
| **1** | 5'-***TT*** | AMP | 10% v/v MPD, 0.040 M sodium cacodylate trihydrate, pH 7.0, 0.012 M spermine tetrahydrochloride, 0.020 M magnesium chloride hexahydrate |
| **2** | 5'-***T***$_m$***C*** | AMP/GMP | 10% v/v MPD, 0.040 M sodium cacodylate trihydrate, pH 7.0, 0.012 M Spermine tetrahydrochloride, 0.08 M sodium chloride, 0.012 M potassium chloride, 0.02 M magnesium chloride hexahydrate |
| **3** | 5'-***T***$_m$***C*** | ApppG | 0.02 M magnesium chloride hexahydrate, 0.05 M MOPS, pH 7.0, 55% v/v Tacsimate, pH 7.0, 2 mM hexammine cobalt(III) chloride |
| **4** | 5'-$_m$***CT*** | ApppG | 0.05 M magnesium sulfate hydrate, 0.05 M HEPES sodium, pH 7.0, 1.6 M lithium sulfate monohydrate |
| **5** | 5'-***CC*** | ApppG | 0.1 M Bis-Tris, pH 6.5, 2 M ammonium sulfate |
| **6** | 5'-***T***$_m$***C*** | Ap-AI-pG | 0.5 M ammonium sulfate, 0.1 M sodium citrate tribasic dihydrate, pH 5.6, 1.0 M lithium sulfate monohydrate |

**Table S2. Data collection statistics.**

| Structure | 1 | 2 | 3 |
|---|---|---|---|
| Space group | P321 | P321 | P321 |
| Unit cell parameters (Å, °) | 43.84, 43.84, 82.84, 90, 90, 120 | 43.53, 43.53, 80.14, 90, 90, 120 | 44.12, 44.12, 84.53 90, 90, 120 |
| Resolution range, Å (last shell) | 50-1.90 (1.97-1.90) | 50-1.45 (1.45-1.5) | 50-1.64 (1.7-1.64) |
| Unique reflections | 7680 | 15931 | 10994 |
| Completeness, % | 100 (100) | 98.5 (100) | 90.1 (100) |
| $R_{merge}$, % | 8.7 (50.1) | 12.2 (53.1) | 13.6 (79.6) |
| $<I/\sigma(I)>$ | 28.9 (3.5) | 12.1 (7.9) | 10.8 (8.1) |
| Redundancy | 14.1 (9.1) | 10.7 (10.8) | 10.1 (10.4) |

| Structure | 4 | 5 | 6 |
|---|---|---|---|
| Space group | P321 | P321 | P3 |
| Unit cell parameters (Å, °) | 44.52, 44.52, 85.52, 90, 90, 120 | 43.54, 43.54, 84.27, 90, 90, 120 | 43.66, 43.66, 81.13, 90, 90, 120 |
| Resolution range, Å (last shell) | 50-1.95 (2.02-1.95) | 50-1.5 (1.55-1.5) | 50-1.76 (1.82-1.76) |
| Unique reflections | 6922 | 15337 | 15744 |
| Completeness, % | 100 (90.8) | 99.8 (98.8) | 91.9 (100) |
| $R_{merge}$, % | 13.8 (41.3) | 8.4 (56.4) | 9.9 (57.7) |
| $<I/\sigma(I)>$ | 10 (13.5) | 21.2 (3.7) | 19.6 (2.3) |
| Redundancy | 8.9 (9.4) | 10.2 (8) | 5.5 (5.5) |

**Table S3. Structure refinement statistics.**

| **Structure** | 1 | 2 | 3 |
|---|---|---|---|
| PDB code | 8SXL | 8SX6 | 8SWO |
| RNA duplex per asymmetric unit | 1 | 1 | 1 |
| Resolution range, Å | 38-1.9 | 26.7-1.45 | 28.19-1.64 |
| $R_{work}$, % | 21.7 | 22 | 24.8 |
| $R_{free}$, % | 26.1 | 24.2 | 24.9 |
| Number of reflections | 7205 | 15101 | 10409 |
| Bond length R.M.S. (Å) | 0.009 | 0.028 | 0.023 |
| Bond angle R.M.S. | 1.794 | 3.611 | 3.845 |
| Average B-factors, (Å$^2$) | 30.9 | 27.9 | 35.9 |

| **Structure** | 4 | 5 | 6 |
|---|---|---|---|
| PDB code | 8SX5 | 8SWG | 8SY1 |
| RNA duplex per asymmetric unit | 1 | 1 | 2 |
| Resolution range, Å | 28.52-1.95 | 37.74-1.5 | 34.29-1.76 |
| $R_{work}$, % | 28.4 | 21.8 | 22.4 |
| $R_{free}$, % | 32.3 | 26.6 | 25.6 |
| Number of reflections | 6597 | 14486 | 14945 |
| Bond length R.M.S. (Å) | 0.019 | 0.021 | 0.022 |
| Bond angle R.M.S. | 3.688 | 3.254 | 2.837 |
| Average B-factors, (Å$^2$) | 26.94 | 27.18 | 25.78 |


[1] L. Li, N. Prywes, C. P. Tam, D. K. O'flaherty, V. S. Lelyveld, E. C. Izgu, A. Pal, J. W. Szostak, *J. Am. Chem. Soc.* **2017**, *139*, 1810-1813.

[2] G. N. Murshudov, P. Skubák, A. A. Lebedev, N. S. Pannu, R. A. Steiner, R. A. Nicholls, M. D. Winn, F. Long, A. A. Vagin, *Acta Crystallogr. D* **2011**, *67*, 355-367.

[3] A. W. Schüttelkopf, D. M. Van Aalten, *Acta Crystallogr. D* **2004**, *60*, 1355-1363.
